# Organellar-specific ROS dynamics drive differential cross-compartmental responses between chloroplast and mitochondria in *C. reinhardtii*

**DOI:** 10.64898/2025.12.07.692803

**Authors:** Gunjan Dhawan, Basuthkar Jagadeeshwar Rao

**Author notes:** Correspondence: Basuthkar Jagadeeshwar Rao.

## Abstract

Reactive oxygen species (ROS) act as key signaling intermediates in plant metabolism, defense, and stress adaptation. In photosynthetic organisms, chloroplast and mitochondria serve as the major hubs of ROS production, thereby coordinating stress responses across cellular compartments. However, the extent of cross-organellar communication following compartmentalized oxidative stress remains poorly understood. Using *C. reinhardtii* as a model system, we induced compartmentalized oxidative stress in chloroplast and mitochondria to investigate whether the ROS generation in one organelle triggers any functional response in the other organelle. Methyl viologen (MV) was used to induce ROS production in the chloroplast, while menadione (MD) was used to trigger mitochondrial ROS production. Real-time monitoring of compartment-specific roGFP strains showed specific, localized, and reversible ROS production following MV and MD treatment as a function of time within the respective target organelles. Comprehensive functional analyses following compartmentalized ROS perturbations revealed that the mitochondrial ROS remained localized to their site of origin and did not detectably affect chloroplast function, as evidenced by the unchanged chlorophyll a fluorescence parameters and photosystem stoichiometry. In contrast, chloroplast-derived ROS diffused from its site of origin and transiently suppressed the mitochondrial oxygen consumption rates. Together, these findings underscore a dynamically regulated system, where chloroplast ROS leaks out and affects mitochondrial function while the mitochondrial ROS remains localized and does not impact chloroplast function. These compartment-specific ROS responses likely represent an adaptive mechanism for coordinating cellular energy metabolism in response to fluctuating environmental conditions.

## Introduction

Reactive oxygen species (ROS), inevitable by-product of aerobic metabolism, are well established as crucial signaling molecules in plants that regulate various cellular processes including growth, cell cycle and development, programmed cell death, pathogen defense, and systemic signaling (Apel & Hirt 2004; Asada 2006; Noctor et al. 2007; Mittler et al. 2004; Møller et al. 2007).

In photosynthetic organisms, chloroplast and mitochondria are the primary sites of ROS production during excess light stress (Asada, 2006; Estavillo et al., 2011; Exposito-Rodriguezet al., 2017; Mittler, 2017; Waszczak et al., 2018; Shapiguzov et al., 2019; Smirnoff & Arnaud, 2019; Foyer & Hanke, 2022). Chloroplast generates ROS species such as superoxide, H_2_O_2_, and singlet oxygen at PSI and PSII, when the light absorption exceeds the photosynthetic capacity (Miller et al., 2010; Asada, 2006; Karuppanapandian et al., 2011). Mitochondria, on the other hand, produce ROS, including H₂O₂ and O_2_^.−^, through mitochondrial electron transport chain components Complex I and Complex III (Navrot et al., 2007; Møller et al. 2007; Noctor et al., 2007). The optimal level of ROS is important during the beneficial interactions of photosynthetic carbon assimilation and mitochondrial metabolism (Dinakar et al., 2010). In plants, both chloroplast and mitochondria undergo several mechanisms, including ROS scavenging systems and retrograde signaling, to maintain the cellular redox balance in response to environmental and metabolic changes. Although both the compartments produce ROS, however, oxidation occurs with different dynamics and amplitudes in these subcellular compartments (Schwarzländer et al., 2009; Rosenwasser et al., 2011; Bratt et al., 2016). The generation of ROS and antioxidant systems within each organelle are characterized but the extent to which ROS generation in one organelle influences the function of another organelle remains poorly understood.

The compartment-specific ROS dynamics can be assessed by inducing oxidative stress conditions that can selectively generate ROS within specific organelles. The physiological impact of perturbing one organelle on another can be transient or permanent, depending on the nature and extent of the perturbation. These physiological changes can affect both photosynthetic and mitochondrial functions, altering energy production, metabolic pathways and cellular functions altogether. Various chemical inducers have been known in the literature that can selectively generate ROS in specific compartments. For instance, methyl viologen (MV) induces ROS in chloroplast by catalyzing the shuttling of electrons from PSI to O₂, resulting in immediate production of O_2_^.−^ (Donahue et al., 1997; Scarpeci et al., 2008; Li et al., 2013; Hawkes, 2014). While menadione (MD) induces ROS production primarily in mitochondria through redox cycling at the electron transport chain (Reichheld et al. 1999; Sweetlove et al., 2002; Senthil-Kumar et al. 2003; Obata et al., 2011; Shi et al., 1994; Monks et al., 1992).

By exploiting these well-characterized compartment-specific ROS inducers, our study aims to understand whether the transient ROS generation within one organelle can influence the function of another subcellular compartment in any manner. We used *Chlamydomonas reinhardtii* as the model system for this study. *C. reinhardti* offers the advantage of perturbing the chloroplast compartment for the activity which is difficult in case of plants (Levine, 1960). Previous reports using roGFP-based sensors have demonstrated a fast and reversible oxidation in response to external H_2_O_2_ (Meyer et al., 2007; Morgan et al., 2011). We used the compartment-specific roGFP (reduction-oxidaton sensitive green fluorescent protein) assay system to monitor the real time ROS dynamics in live cells following MV and MD treatment. After validating the compartmentalized response of MV and MD, we performed the functional analyses, where we observed differentially localized ROS responses at sublethal drug concentrations. The compartmentalized ROS in mitochondria did not impact chloroplast function in any discernible manner. In contrast, chloroplast ROS diffused transiently, suppressing the mitochondrial respiration in a time-dependent manner, thus leading to inter-compartmental effect. However, at higher concentrations cellular lethality was observed. These results suggest that during mild oxidative stress, each organelle possesses specialized mechanisms to counteract excess ROS and thereby maintain cellular homeostasis dynamically.

## Materials and methods

### Strains, media, and growth condition

*Chlamydomonas reinhardtii* wild-type cc-125 strain was obtained from the Chlamydomonas resource center (https://www.chlamycollection.org). The compartment-specific, genetically encoded fluorescent H₂O₂ sensor for Chlamydomonas were kindly provided by Prof. Michael Schroda (Molecular biotechnology & systems biology, Technical University of Kaiserslautern) (Niemeyer et al., 2021). These roGFP sensors were generated in the UVM4 expression strain background (Neupert et al., 2009). *C. reinhardtii* cells were grown under continuous light illumination at a light intensity of 60 µmol m⁻² s⁻ ¹ at 24⁰C. The cells were grown in Tris phosphate (TP) liquid medium with continuous shaking as described (Harris, 2009).

For the compartment-specific ROS production, methyl viologen (Sigma 856177) and menadione (Sigma M5750) were used.

### Chlorophyll Bleaching and cell viability assay

Wild type cc-125 cells were grown photoautotrophically to the mid log phase. The cells were exposed to different concentrations of ROS-inducing agents MV and MD. Briefly, cultures containing equal number of cells (1 × 10^6^ cells/ml) treated with different concentrations of MV (0 µM, 0.2 µM, 0.25 µM, 0.3 µM, 0.5 µM, and 1 µM) and MD (0 µM, 10 µM, 20 µM, 40 µM, 50 µM, and 100 µM) were photographed at 24 hours to get the chlorophyll pigmentation levels. A decrease in chlorophyll content, often visualized as a loss of green pigmentation (bleaching) indicated oxidative damage to the chloroplast and a potential defect in photosynthetic function (Doi et al., 2001; Allorent et al., 2013; Hemschemeier, 2013; Ugalde et al., 2021). For the cell viability assay, cells treated with different concentrations of MV and MD were stained with trypan blue, such that the dead cells retained the blue stain due to loss of membrane integrity, allowing a clear distinction from the live cells (Strober, 2015). The percentage of live cells relative to the total cell count was plotted after treatment with MV or MD across different time points.

### Confocal imaging for the compartment-specific ROS detection

The compartment-specific genetically encoded, fluorescent H₂O₂ sensors for *Chlamydomonas* were employed for the detection of ROS, using live cell confocal imaging. Specifically, chloroplast (pAR-CDJ1-roGFP2-Tsa2ΔCR-MS) and mitochondrial (pAR-mtTP70C-roGFP2-Tsa2ΔCR-MS) roGFP strains were treated with methyl viologen (0 µM, 0.2 µM, and 0.25 µM MV) or menadione (0 µM, 20 µM, and 40µM) and imaged for the GFP fluorescence at different time intervals. Briefly, the cells (1×10⁶ cells/ml) were immobilized on a 0.8% low melting agarose slide followed by imaging for roGFP (excitation 488 nm; emission 518-574 nm) and autofluorescence of chloroplast (excitation 561 nm; emission 685-750 nm) using appropriate filters in Leica TCS SP8 confocal laser scanning microscope. The images were obtained as Z-stacks and processed with Fiji Software.

Chloroplast ROS showed three different patterns - normal (basal ROS), punctate (intermediate ROS), and diffused (high ROS, accompanied by the visible distortions in the chloroplast morphology). Similarly, mitochondrial ROS showed these three patterns - normal, intermediate, and high. Each cell was assessed for the dominant pattern of ROS and the frequency distribution of cells exhibiting these patterns was quantified for the untreated (control), MV and MD treated samples at different time points. The data from 50 cells was statistically analyzed, quantified using ImageJ software and plotted using GraphPad Prism 9.5.1 software.

### Handy-PEA (photosynthetic efficiency analyzer)

Handy-PEA (a portable Photosynthetic Efficiency Analyzer) by Hansatech Instr. Ltd, Kings Lynn, Norfolk, UK was used to measure OJIP fluorescence transients of MD-treated samples (0 µM, 20 µM, and 50 µM MD) at 0.5 hr, 1 hr, 3 hr, 6 hr, and 12 hr. The OJIP transients were recorded for 1 s with an excitation light wavelength of 650 nm and 3000 μmol photons m^−2^ s^−1^ light intensity in liquid cell cultures. The “OJIP” stands for the four phases of the transient: O (initial fluorescence level), J (fluorescence increase to peak), I (inflection point), and P (peak fluorescence). These phases provide insights into the functional state of photosystem II and can be indicative of stress or the electron transport or carrier activity (Stirbet & Govindjee, 2011; Kalaji et al., 2012; Tsimilli-Michael, 2020). Briefly, 1ml (1×10^6^ UVM4 cells/ml) of dark incubated liquid cultures were taken in a cuvette and OJIP transients were recorded. The maximum quantum efficiency of PSII (Fv/Fm) was calculated using the following formula Fv/Fm = (Fm – Fo)/Fm. Where Fv is the variable fluorescence, Fm is the maximum fluorescence, and Fo is the minimum fluorescence recorded at 50μs after the onset of illumination.

### 77K Spectroscopy

The low temperature (77K) facilitates the trapping of photosynthetic complexes in a specific state, enabling detailed spectroscopic analyses. The fluorescence emission spectrum at 77K was recorded using a Jasco FP8500 spectrophotometer. A special dewar flask made of quartz was used to measure the fluorescence at very low temperature (liquid nitrogen temperature). The parameters were set as follows: excitation wavelength 440 nm; Emission bandwidth 660-760 nm; excitation and emission slit width 5 nm each. UVM4 cells treated with MV at a concentration of 0 µM, 0.2 µM, and 1 µM were pelleted at 3hr, 6hr, 12hr, 24hr, and 48hr. The pellets were resuspended in 500 µl TP media, and the cells were immediately taken for fluorescence measurement in a liquid nitrogen dewar flask (Cardol et al., 2003; Snellenburg et al., 2017). The emission traces were obtained with noise and the final spectral traces were normalized and plotted using GraphPad prism 9.5.1 Software.

### Seahorse XFp Flux Analyzer assay

MV-treated UVM4 cells (2×10⁶) at different timepoints (3hr, 6hr, 12hr, 24hr, and 48hr) were pelleted and resuspended in fresh TP media. For the Sea horse mitochondrial flux assay, these cells were pipette into each well in the 8-well flux analyzer plate, followed by measurement of oxygen consumption rate (OCR) as per the Agilent Guidelines (https://www.agilent.com/cs/library/usermanuals/public/XF_Cell_Mito_Stress_Test_Kit_User_G <u>uide.pdf</u>). Two of the wells were left blank for blank subtraction. CCCP (1 µM) (Sigma C2759-100 mg) was injected in Port A of each well of the seahorse cartridge. Each measurement comprised of a calibration cycle of 40 minutes followed by an equilibration cycle of 12 minutes, where the cells were mixed and incubated in the dark for 12 minutes, and then the OCR measurements were recorded at 24⁰C. OCR measurement consisted of a 1 minute mix cycle (to oxygenate the microchamber), a 1 minute wait period (allow settling of cells), and a 3 minute of measurement period (https://www.agilent.com/cs/library/usermanuals/public/S7894-10000_Rev_B_Wave_2_4_User_Guide.pdf). 3 OCR readings were taken for the basal and maximal respiration each. Multiple technical and biological replicates were taken to obtain reliable and reproducible data. Statistical analyses and the graph were plotted using GraphPad Prism 9.5.1 Software.

### Mitochondrial morphology changes

Live cell confocal imaging was performed using mitochondrial GFP strain (MDH4-GFP) for the visualization of mitochondrial morphologies of *C. reinhardtii* cells following MV and MD treatment. Briefly, the cells (1×10⁶ cells/ml) were immobilized using 0.8% low melting agarose on a coverslip, followed by imaging for mitochondrial GFP (excitation 488 nm; emission 518-574 nm) and autofluorescence of chloroplast (excitation 561 nm; emission 685-750 nm). The images were obtained as Z-stacks and processed with Fiji Software. Mitochondrial morphology was classified as tubular, intermediate, and fragmented, as per the published protocol (Rambold et al., 2011). Each cell was assessed for the mitochondrial phenotype (dominant), and the frequency distribution of cells exhibiting these phenotypes was quantified. The data from ∼50 cells was statistically analyzed, quantified using ImageJ software, and plotted using GraphPad Prism 9.5.1 software.

### Statistical analysis

All the statistical analyses, including arithmetic mean, standard deviation (S.D), and wherever applicable 2-way anova was performed in all cases. 2 way anova ordinary 2 data sets with Bonferroni test was performed using GraphPad Prism version 9.5.1 for windows, GraphPad Software (www.graphpad.com).

## Results

### Transient and localized ROS production by methyl viologen (MV) and menadione (MD)

Methyl viologen (MV) primarily produces ROS in the chloroplast, while menadione (MD) triggers ROS production in the mitochondrial compartment (Donahue et al., 1997; Sweetlove et al., 2002; Scarpeci et al., 2008; Obata et al., 2011; Li et al., 2013; Hawkes, 2014). In order to establish the sublethal concentrations of MV and MD, that can perturb redox homeostasis without compromising the cell survival, we performed chlorophyll bleaching and cell viability assays. We observed that cells treated with MV at concentrations ≤ 0.25µM and MD at concentrations ≤ 40µM remained viable and therefore can be used to analyze the ROS dynamics further (**Figure 1**). Beyond these concentration ranges, cell death was observed as a function of time. These results are consistent with the previously reported concentration range of MV (Fischer et al., 2010; Jamers & Coen, 2010) and MD (Sirisha et al., 2014; Khona et al., 2016; Marinovska et al., 2022). After determining the sublethal concentrations, we examined the compartment specificity of these ROS inducers.

**Figure 1:**
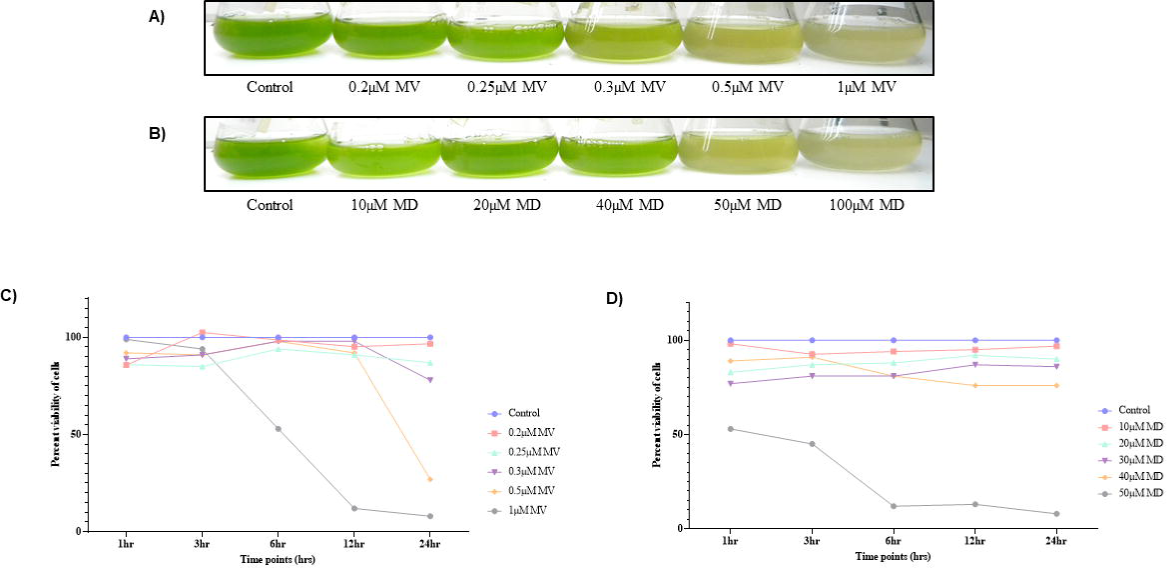
Chlorophyll bleaching and cell viability assay to determine methyl viologen (MV) and menadione (MD) concentration. **(A)** Photoautotrophic *C. reinhardtii* wild-type cc125 cells treated with Methyl viologen concentration - 0 (control), 0.2 µM MV, 0.25 µM MV, 0.3 µM MV, 0.5 µM MV, and µM MV were photographed at 24hr time point to observe chlorophyll pigmentation. **(B)** *C. reinhardtii* cells treated with Menadione concentration – 0 (control), 10 µM MD, 20 µM MD, 40 µM MD, 50 µM MD, and 100 µM MD were photographed at 24 hr time point for chlorophyll pigmentation. The percentage of viable cells at different concentrations of methyl viologen **(C)** and menadione **(D)** at 1hr, 3hr, 6hr, 12hr, and 24hr were quantified using hemocytometer and plotted using GraphPad Prism 9.5.1

We utilized roGFP-based redox sensing system to assess the impact of MV and MD on live cells. roGFP strains were obtained, which were generated using Chlamydomonas Modular Cloning toolbox to express a GFP-tagged hypersensitive H_2_O_2_ sensor (Tsa2ΔCr). This sensor device was targeted to specific compartments, including cytosol, nucleus, mitochondria, and chloroplast, using specific promoters (Niemeyer et al., 2021).

Using live-cell confocal imaging, we visualized ROS production in the roGFP strains targeted to the chloroplast (pAR-CDJ1-roGFP2-Tsa2ΔCR-MS) and mitochondria (pAR-mtTP70C-roGFP2-Tsa2ΔCR-MS) following MV and MD treatment (**Figure 2a and 3a**, respectively). As expected, we observed that MV induces ROS generation specifically in the chloroplast compartment (**Figure 2 and Table S1 and S4**), whereas MD predominantly induces ROS production in mitochondria (**Figure 3 and Table S2 and S3)**. However, the specificity of these ROS inducers in the respective compartment is time-dependent and dynamic, suggesting that ROS levels are modulated in a transient manner as a function of time in the current assay system. So, we asked whether this transient induction of ROS in one compartment impacts the function of the other compartment in any manner.

**Figure 2:**
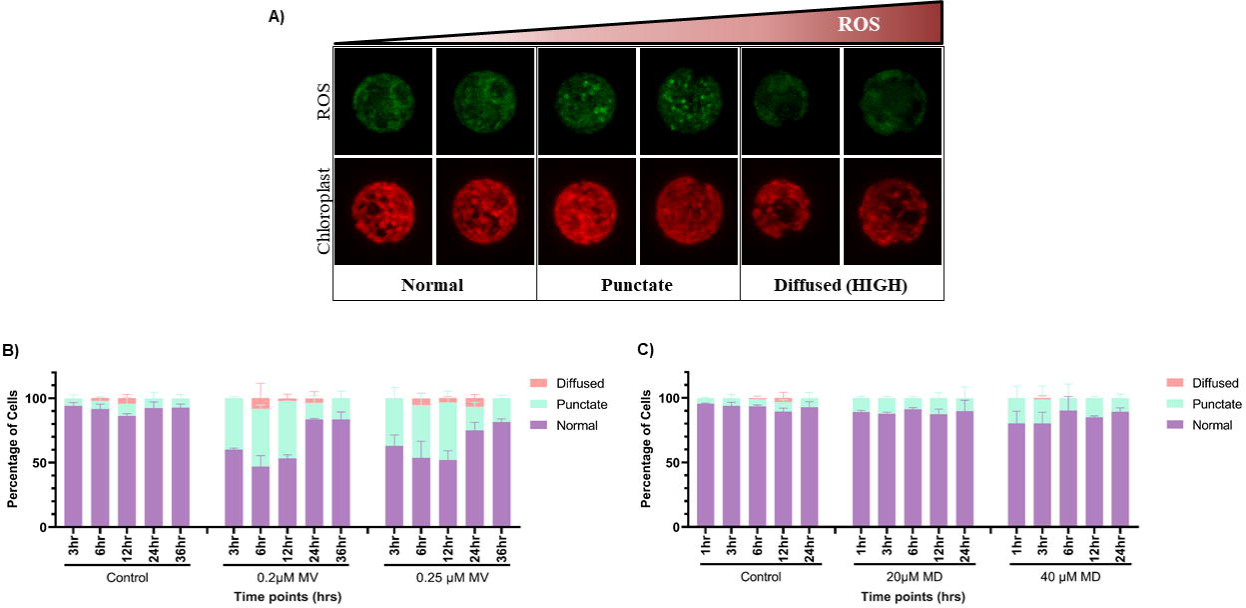
Distribution of chloroplast-specific roGFP signal in *C. reinhardtii* cells upon MV and MD treatment. *C.reinhardtii* chloroplast-specific roGFP strain (pAR-CDJ1-roGFP2-Tsa2ΔCR-MS) was treated with different concentrations of methyl viologen and menadione separately and imaged using Leica TCS SP8 confocal laser scanning microscope. **(A)** Representative images of roGFP signal across ROS gradient in chloroplast compartment. Three patterns were obtained depending on the ROS gradient – basal ROS (labelled as ‘Normal’), intermediate level of ROS (labelled as ‘Punctate’), and high level of ROS (labelled as ‘Diffused’). **(B)** The percentage of cells having these 3 ROS patterns at different timepoints (3hr, 6hr, 12hr, 24hr, 36hr, and 48hr) upon Methyl viologen treatment at 0µM, 0.2µM, and 0.25µM MV concentration was quantified. **(C)** Similarly, quantification of percentage of cells having these 3 ROS patterns at different timepoints (1hr, 3hr, 6hr, 12hr, and 24hr) upon Menadione treatment at 0µM, 20µM, and 40µM MD concentration was performed. The data represents an average of two independent biological repeats. Error bars represent Standard Deviation (SD). Statistical analyses were performed using 2-way ANOVA.

**Figure 3:**
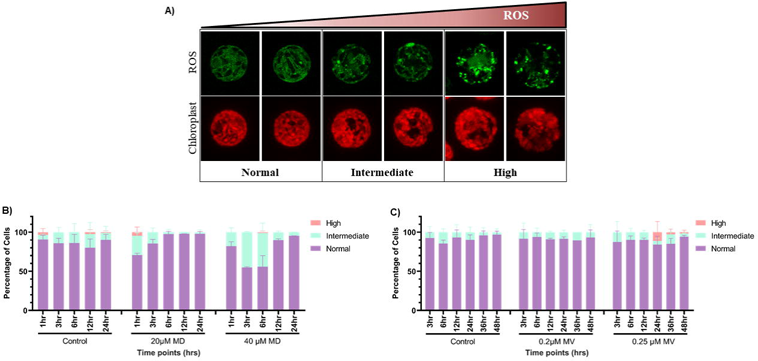
Distribution of mitochondria-specific roGFP signal in *C. reinhardtii* cells upon MD and MV treatment. *C.reinhardtii* mitochondria-specific roGFP strain (pAR-mtTP70C-roGFP2-Tsa2ΔCR-MS) was treated with different concentrations of methyl viologen and menadione separately and imaged using Leica TCS SP8 confocal laser scanning microscope. **(A)** Representative images of roGFP signal across ROS gradient in mitochondrial compartment. Three ROS patterns were obtained depending on the ROS gradient – basal ROS (labelled as ‘Normal’), intermediate level of ROS (labelled as ‘Intermediate’), and high level of ROS (labelled as ‘High’). **(B)** The percentage of cells having these 3 ROS patterns at different timepoints (1hr, 3hr, 6hr, 12hr, and 24hr) upon Menadione treatment at 0µM, 20µM, and 40µM MD concentration was quantified using Image J software and plotted using Graphpad Prism 9.5.1. **(C)** Similarly, quantification of percentage of cells having these 3 ROS phenotypes at different timepoints (3hr, 6hr, 12hr, 24hr, 36hr, and 48hr) upon Methyl viologen treatment at 0µM, 0.2µM, and 0.25µM MV concentration was performed. The data represents an average of two independent biological repeats. Error bars represent Standard Deviation (SD). Statistical analyses were performed using 2-way ANOVA.

### No discernible change in chloroplast activity in response to mitochondrial ROS perturbation

We induced ROS in mitochondrial compartment using MD (0 µM, 20 µM, and 50 µM MD) and assessed its impact on the chloroplast function (using Handy PEA and 77K spectroscopy). Handy PEA is used to measure the chlorophyll a fluorescence, a widely recognized indicator of photosynthetic activity (Hornyák & Płażek., 2024). 77K fluorescence spectroscopy measures the relative abundance of the two photosystems (PSI and PSII) at low temperature (77K) (Minagawa, 2011; Goldschmidt-Clermont & Bassi, 2015). Using these assay systems, we observed no significant change in the OJIP transients, Fv/Fm values, and F (PSI/PSII) levels in the MD-treated samples compared to the control, suggesting that mitochondrial ROS did not impact chloroplast function in any discernible manner (**Figure 4, 5 and Table S5, S6** ). In contrast, at lethal concentration of MD, mitochondrial ROS spills over, impacting the functions of chloroplast, thereby causing cellular cytotoxicity. This shift from a localized response at sublethal concentration to a system-wide response at lethal concentration suggests that ROS mediated inter-organellar communication may emerge only when the mitochondrial antioxidant buffering capacity exceeds a certain threshold. Based on these observations, further we assessed the impact of chloroplast ROS on mitochondrial function.

**Figure 4:**
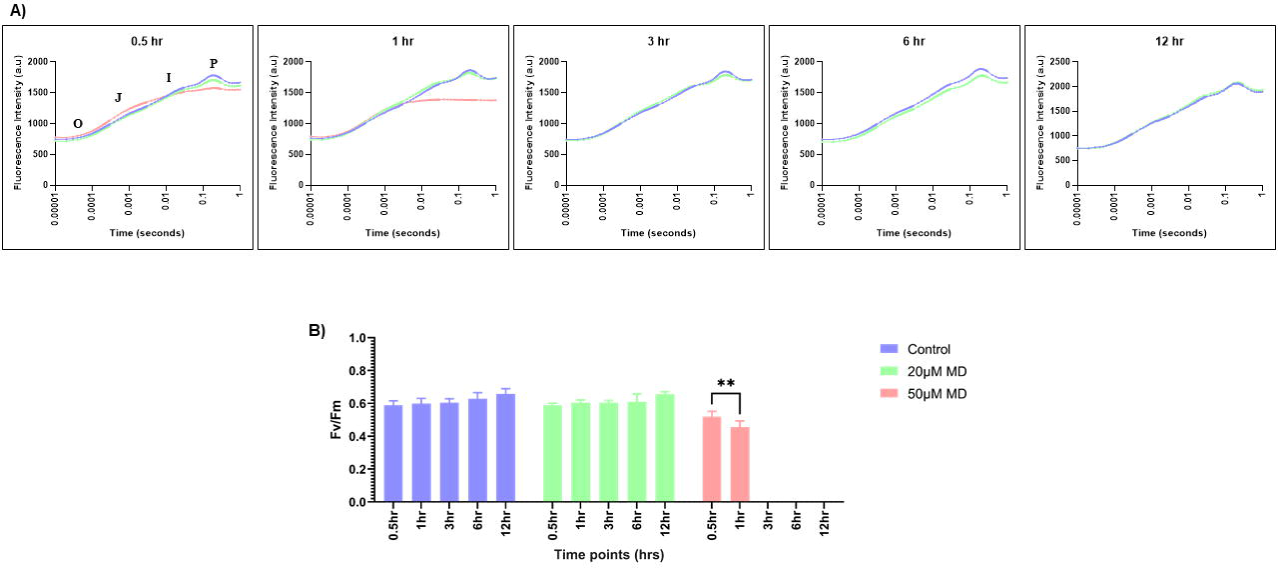
OJIP fluorescence transients were recorded using Handy PEA (a portable Photosynthetic Efficiency Analyzer) for MD-treated cells. Untransformed UVM4 cells were treated with different MD concentrations and OJIP transients were recorded at specified timepoints. **(A)** Average OJIP transients were plotted for all three MD concentrations - 0µM (blue), 20µM (green), and 50µM MD (orange), across all the time points (0.5hr, 1hr, 3hr, 6hr, 12hr, and 24hr). **(B)** The ratio of Fv/Fm for all three conditions 0µMMD (blue), 20µMMD (green) and 40µMMD (orange) at all the time points were plotted. The data represents an average of three independent biological repeats. Error bars represent Standard Deviation (SD). Statistical analyses were performed using 2-way ANOVA.

**Figure 5:**
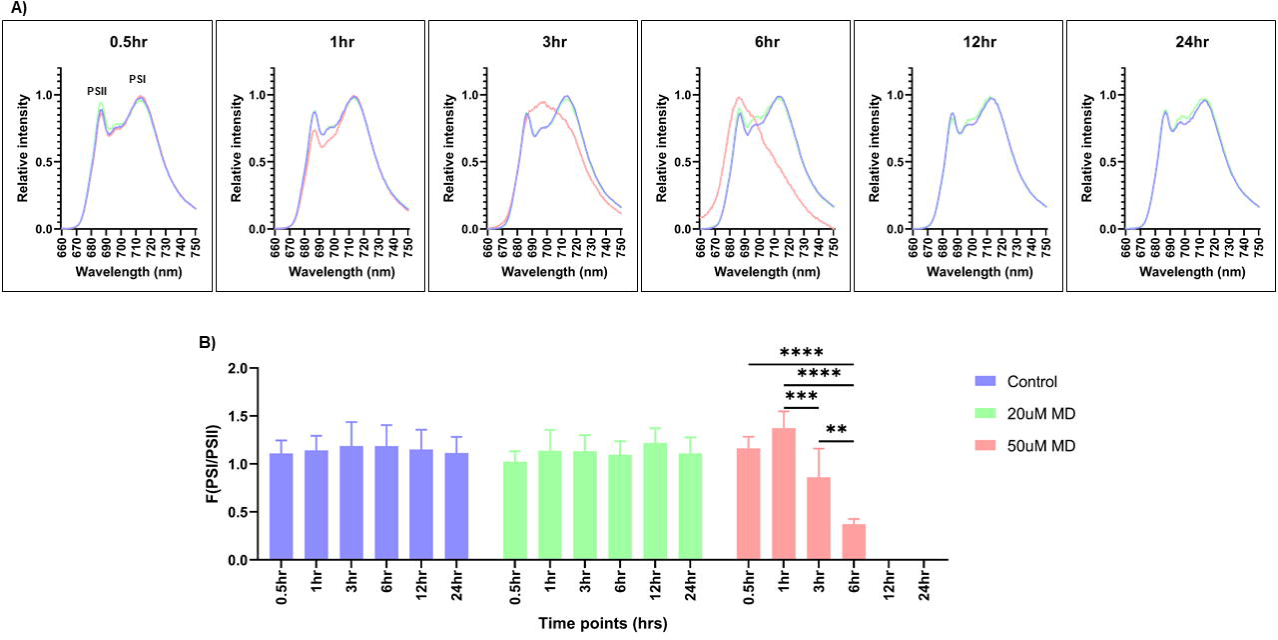
PSI and PSII spectral traces were recorded using 77K fluorescence spectroscopy for MD-treated cells. Untransformed UVM4 cells were treated with different MD concentrations and spectral readings were taken at different timepoints. **(A)** Average fluorescence emission spectral traces (77K spectra) were plotted for all three MD concentrations - 0µM (blue), 20µM (green), and 50µM MD (orange), across all the time points (0.5hr, 1hr, 3hr, 6hr, 12hr, and 24hr). **(B)** The ratio of F(PSI/PSII) for all three conditions 0µMMD (blue), 20µMMD (green) and 40µMMD (orange) at all the time points were plotted. The data represents an average of three independent biological repeats. Error bars represent Standard Deviation (SD). Statistical analyses were performed using 2-way ANOVA.

### Transient change in mitochondrial activity in response to chloroplast ROS

To evaluate the mitochondrial activity changes following MV treatment, we measured mitochondrial respiration rate using Seahorse XFp Flux analyzer. It provides the intracellular readout of oxygen consumption rate [OCR (pmol/min per 1×10^6^ cells)] at 24⁰C in the dark as a function of time (Upadhaya & Rao, 2019; Dhawan & Rao, 2025).

In contrast to the mitochondrial ROS, which showed no change in chloroplast function, chloroplast ROS showed a transient change in the mitochondrial oxygen consumption rate (**Figure 6 and Table S7**). The mitochondrial OCR levels were initially low (∼43) compared to the control cells (∼78-87), suggesting suppressed respiratory activity during early chloroplast oxidative imbalance. However, once the chloroplast ROS balance was restored, mitochondrial OCR levels recovered and attained baseline respiration rate (∼79) similar to the control condition. This dynamic change suggests that ROS diffuses out from the chloroplast and impacts mitochondrial function in a time-dependent manner. In contrast, at lethal concentration of MV, chloroplast ROS spill over uncontrollably, resulting in sustained respiratory impairment and thereby causing cellular cytotoxicity. Together, these results suggest that chloroplast ROS diffuses out and impacts mitochondrial function. Further, we assessed whether these functional changes were accompanied by the mitochondrial morphology changes following MV and MD treatment.

**Figure 6:**
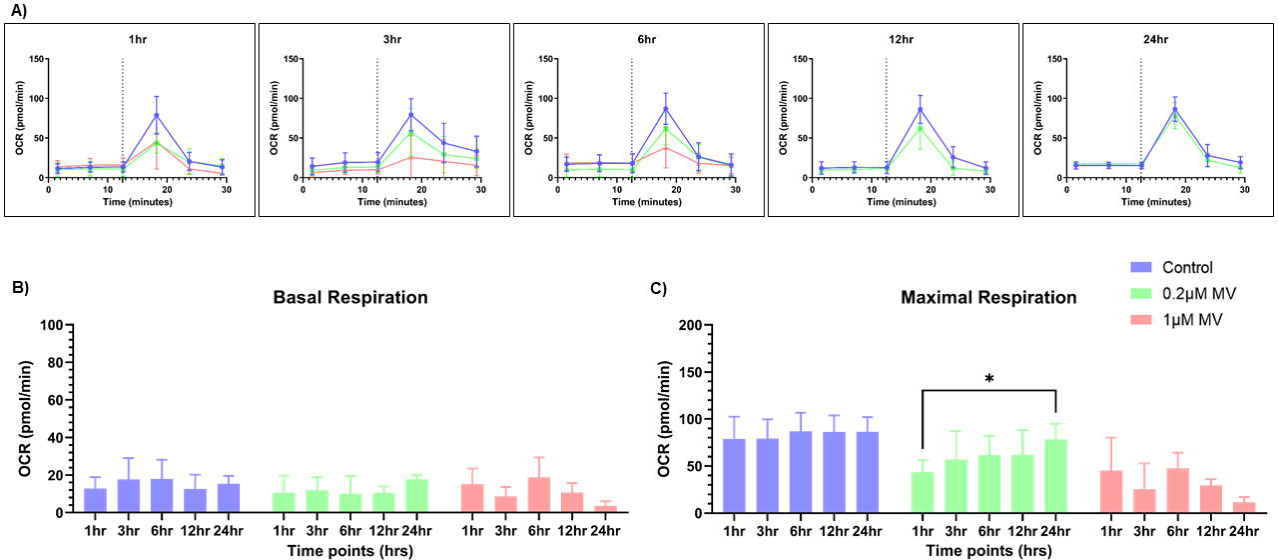
Mitochondrial respiration rate was measured for the MV-treated samples. Untransformed UVM4 cells were treated with different concentrations of MV and OCR readings were taken at different timepoints. **(A)** Seahorse Flux analyzer assay was performed for the oxygen consumption rate (OCR) measurements for all three MV concentrations 0µM (blue), 0.2µM (green), and 1µM (orange) MV at 1hr, 3hr, 6hr, 12hr, 24hr timepoints. The basal respiration **(B)** and maximal respiration after the addition of CCCP **(C)** were plotted for MV-treated samples at 0µM (blue), 0.2µM (green), and 0.25µM (orange) MV across all time points. The data represents an average of three independent biological repeats. Error bars represent Standard Deviation (SD). Statistical analyses were performed using 2-way ANOVA.

### No measurable change in mitochondrial morphology following MV treatment

We imaged the live *C. reinhardtii* cells to observe the mitochondrial morphology changes following MV and MD treatment. We observed that majority of cells were having tubular morphology following MV treatment (**Figure 7**). There was no measurable morphology change in response to MV treatment compared to the control samples (**Figure 7 and Table S8)**. However, a transient morphology change from tubular to fragmented and intermediate forms following MD treatment was observed (**Figure 7 and Table S9)**. The absence of detectable mitochondrial ROS and morphology change following MV treatment, yet the dynamic change in the mitochondrial function suggests that even the undetectable levels of ROS from the chloroplast impact the mitochondria activity. However, this change is transient and time-dependent.

**Figure 7:**
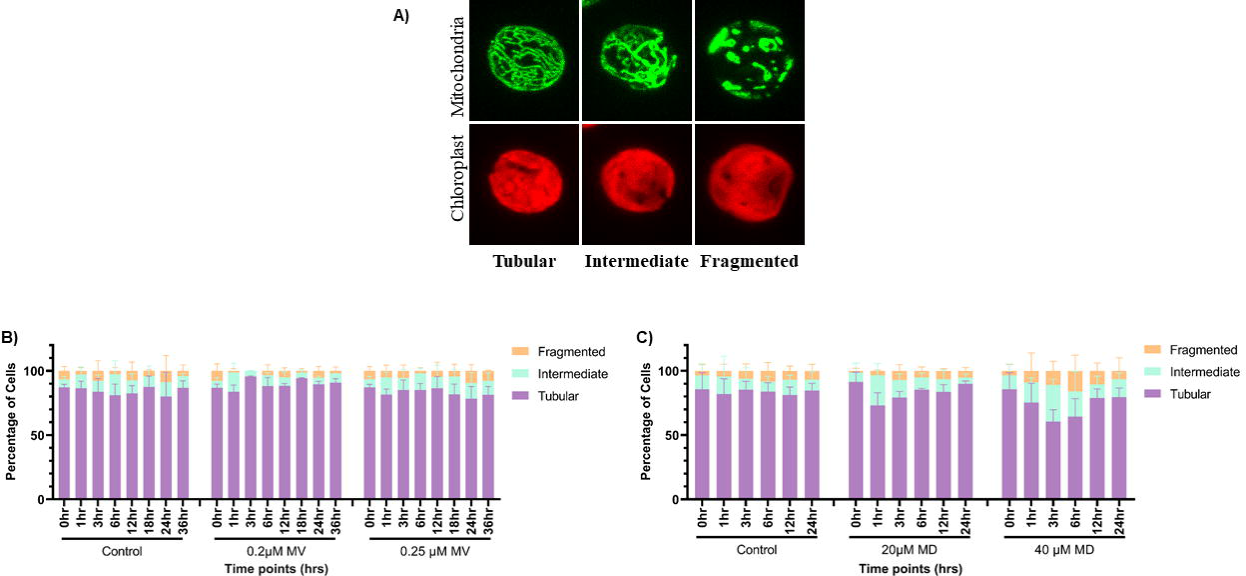
Quantification of cells having different mitochondrial morphologies upon MV and MD treatment. Live cell confocal imaging was performed using photoautotrophic *C. reinhardtii* mitochondrial GFP (MDH4-GFP) cells treated with different concentrations of MV and MD. MDH4-GFP cells were imaged for fluorescence at 488nm to visualize mitochondria. **(A)** Representative images of mitochondrial morphologies obtained upon MV and MD treatment – Tubular, Intermediate, and Fragmented. **(B)** Quantification of the percentage of cells having these 3 mitochondrial morphologies at different timepoints (0hr, 1hr, 3hr, 6hr, 12hr, 24hr, 36hr, and 48hr) upon Methyl viologen treatment at 0µM, 0.2µM and 0.25µM MV concentration was plotted. **(C)** The percentage of cells having different mitochondrial morphologies at specified timepoints (0hr, 1hr, 3hr, 6hr, 12hr, and 24hr) upon Menadione treatment at 0µM, 20µM, and 40µM MD concentration was quantified and plotted. The data represents an average of three independent biological repeats. Error bars represent Standard Deviation (SD). Statistical analyses were performed using 2-way ANOVA.

## Discussion

In the present study, we tried to address a fundamental question in the cellular redox biology: whether transient ROS generation within one organelle can influence the function of another subcellular compartment in any manner. Using organelle-specific ROS inducers, our findings demonstrate for the first time in *C. reinhardtii* that highly compartmentalized ROS in mitochondria remains localized and does not detectably impact chloroplast function whereas, in the same system, chloroplast ROS exhibits the capacity to diffuse out of the site of origin and impact the mitochondrial function in a transient manner. The observed compartmentalized redox responses can be explained by the fundamental differences in the ROS metabolism and detoxification mechanisms in the two organelles.

Mitochondrial ROS, primarily generated at the electron transport chain complexes I and III, are generally short-lived and rapidly detoxified by the matrix-localized antioxidant systems, including manganese superoxide dismutase (MnSOD), glutathione peroxidase, and peroxiredoxins (Morgan et al., 2008; Lázaro et al., 2013). This detoxification ensures that mitochondrial ROS remain largely confined within the organelle and does not diffuse to other compartments, maintaining localized redox homeostasis. In contrast, chloroplast redox imbalance is known to generate diffusible ROS, such as H_2_O_2_ or singlet oxygen, which are capable of diffusing beyond the site of origin and thereby triggering retrograde signaling pathways and short term redox adjustments in mitochondria or cytosol (Fischer *et al*. 2007; Mullineaux & Karpinski 2002; Baier & Dietz 2005; Nott *et al*. 2006; Mullineaux 2009; Pfannschmidt *et al*. 2009). The transient change in the mitochondrial respiration following chloroplast ROS production observed in our study compliments the previous reports showing that chloroplast derived ROS can modulate mitochondrial energy metabolism and electron transport fluxes under stress conditions (Raghavendra & Padmasree 2003; Dinakar *et al*. 2010; Yoshida & Noguchi, 2011). Together these findings suggest real-time metabolic coordination between the chloroplast and mitochondria to maintain cellular homeostasis.

The differential inter-organellar ROS responses observed in our study have important implications on how photosynthetic organisms maintain energy homeostasis under fluctuating environmental conditions. The photosynthetic organisms frequently experience transient oxidative imbalances that mimic the compartment-specific ROS perturbations induced in our study. For instance, physiological conditions such as fluctuating light, nutrient limitation, and temperature stress naturally generate transient and localized ROS responses in chloroplast and mitochondria, rather than just causing cytotoxic effects. High light or fluctuating light conditions enhance chloroplast ROS formation through over-reduction of photosynthetic ETC, which can be correlated with expression of antioxidant and defense genes, and phosphorylation of thylakoid proteins (Dat *et al*. 2000; Vranová *et al*. 2002; Karpinski *et al*. 1999; Zer & Ohad 2003; Mühlenbock *et al*. 2008; Li *et al*. 2009). Acclimation to changes in light quality and intensity requires balancing energy distribution within the photosynthetic machinery to efficiently drive photosynthesis and to protect the thylakoid from damage.

On the other hand, mitochondrial respiration plays a complementary role in neutralizing excess reducing equivalents, thereby preventing oxidative damage to thylakoid membranes and other cellular components (Møller 2001; Raghavendra & Padmasree 2003; Dinakar *et al*. 2010). However, mitochondrial function during increased respiratory and photorespiratory activity is sensitive to oxidative damage, similar to the damage induced by methyl viologen, chilling or drought (Suzuki et al, 2012). Oxidized lipids such as polyunsaturated fatty acids generated under these conditions inhibit tricarboxylic acid (TCA) cycle activity, causing perturbation in carbon and nitrogen metabolisms (Taylor et al., 2002, 2004; Mueller 2004; Møller *et al*. 2007; Ito *et al*. 2009). These ROS are buffered by antioxidant systems such as thioredoxin-peroxiredoxin, thus connecting ROS to the reversible redox signaling rather than immediate cytotoxicity. The resulting ROS in these conditions facilitates immediate stress response as well as long-term acclimation to stress, thereby enhancing plant resilience and survival in fluctuating environmental conditions.

Additionally, the transient nature of mitochondrial response to chloroplast ROS suggests the existence of feedback mechanisms that restore baseline respiration once the chloroplast redox balance is restored, preventing chronic disruption of cellular energy balance. Therefore, understanding these inter-organellar ROS responses is crucial for comprehending how photosynthetic organisms acclimate to the changing environmental conditions and may help in developing the strategies to improve stress tolerance.

## Conclusion

Our study highlights the differential inter-organellar ROS communication between chloroplast and mitochondria in a dynamically regulated system. Chloroplast-derived ROS diffuse transiently affecting mitochondrial function in a time-dependent manner, while the mitochondrial ROS remains compartmentalized without affecting chloroplast functions. These differential ROS responses reflect differences in ROS metabolism and detoxification mechanism between chloroplast and mitochondria. The transient and reversible nature of these ROS responses at sublethal concentrations of MV and MD underscores an adaptive mechanism that enables photosynthetic organisms to maintain cellular homoeostasis under fluctuating environmental conditions. Further investigation using the organellar-specific partial loss-of-function mutants is required to test for cross-regulation of activity under natural environmental fluctuations to fully elucidate the physiological mechanisms underlying these inter-organellar ROS responses.

## Data Availability Statement

The raw data supporting the conclusions of this article will be made available by the authors, without undue reservation.

## CRediT authorship contribution statement

**G.D:** Writing – original draft, Conceptualization, Data curation, Formal analysis, Investigation, Methodology, Resources, Software, Validation, Visualization, Writing- review and editing.

**B.J.R:** Conceptualization, Funding acquisition, Project administration, Resources, Supervision, Writing- review and editing.

## Acknowledgments

We acknowledge the assistance from IISER Tirupati and University of Hyderabad faculty and staff for carrying out the experiments in this study. We express our sincere gratitude to Prof. Michael Schroda (Technical University of Kaiserslautern, Kaiserslautern, Germany) and Prof. Ralph Bock (Max Planck Institute of Molecular Plant physiology, Potsdam, Germany) for providing the compartment-specific roGFP strains and the untransformed background expression strain, UVM4. A special thanks to Prof. N Prakash Prabhu, UoH and Prof. Naresh Babu Sepuri, UoH, for providing the 77K Spectroscopy and Sea Horse facility in the department. We acknowledge JC Bose fellowship grant (DST) [10X-217 to B.J. Rao], MHRD intramural funding to IISER Tirupati to B.J. Rao; University of Hyderabad funding to B.J. Rao.

## Abbreviations

ROS: Reactive oxygen species
MV: Methyl viologen
MD: Menadione
roGFP: reduction-oxidaton sensitive green fluorescent protein
PSI and PSII: Photosystem I and II
H2O2: Hyderogen peroxide
O_2_^.−^: Superoxide
TP: Tris phosphate
OCR: Oxygen consumption rate
MnSOD: Manganese superoxide dismutase
ETC: Electron transport chain
TCA: Tricarboxylic acid cycle.

## Supplementary Material

**Supplementary Table 1.** Quantification of the percentage of cells having the 3 ROS phenotypes in the chloroplast compartment at different timepoints (3hr, 6hr, 12hr, 24hr, 36hr, and 48hr) upon Methyl viologen treatment at 0µM, 0.2µM, and 0.25µM MV concentration. All values represent mean ± SD, n=2.

**Supplementary Table 2.** Quantification of percentage of cells having the 3 ROS phenotypes in the chloroplast compartment at different timepoints (1hr, 3hr, 6hr, 12hr, and 24hr) upon Menadione treatment at 0µM, 20µM, and 40µM MD concentration. All values represent mean ± SD, n=2.

**Supplementary Table 3.** Quantification of the percentage of cells having the 3 ROS phenotypes in the mitochondrial compartment at different timepoints (1hr, 3hr, 6hr, 12hr, and 24hr) upon Menadione treatment at 0µM, 20µM, and 40µM MD concentration. All values represent mean ± SD, n=2.

**Supplementary Table 4.** Quantification of percentage of cells having the 3 ROS phenotypes in the mitochondrial compartment at different timepoints (3hr, 6hr, 12hr, 24hr, 36hr, and 48hr) upon Methyl viologen treatment at 0µM, 0.2µM, and 0.25µM MV concentration. All values represent mean ± SD, n=2.

**Supplementary Table 5.** Quantification of the normalized ratio of Fv/Fm for all three conditions 0µM MD, 20µM MD, and 50µM MD at all the time points. All values represent mean ± SD, n=3.

**Supplementary Table 6.** Quantification of the ratio of F (PSI/PSII) for all three conditions 0µM MD, 20µM MD, and 50µM MD at all the time points. All values represent mean ± SD, n=3.

**Supplementary Table 7.** The basal respiration **(A)**, and maximal respiration rates **(B)** were quantified for MV-treated samples at 0µM, 0.2µM, and 1µM MV across all time points. All values represent mean ± SD, n=3.

**Supplementary Table 8.** Quantification of the percentage of cells having these 3 mitochondrial morphologies at different timepoints (0hr, 1hr, 3hr, 6hr, 12hr, 24hr, 36hr, and 48hr) upon Methyl viologen treatment at 0µM, 0.2µM and 0.25µM MV concentration. All values represent mean ± SD, n=3.

**Supplementary Table 9.** Quantification of the percentage of cells having different mitochondrial morphologies at specified timepoints (0hr, 1hr, 3hr, 6hr, 12hr, and 24hr) upon Menadione treatment at 0µM, 20µM, and 40µM MD concentration. All values represent mean ± SD, n=3.

