## Supplementary data for "Organellar-specific ROS dynamics drive differential cross-compartmental responses between chloroplast and mitochondria in *C. reinhardtii*"

**Table 1.**

Quantification of the percentage of cells having the 3 ROS phenotypes in the chloroplast compartment at different timepoints (3hr, 6hr, 12hr, 24hr, 36hr, and 48hr) upon Methyl viologen treatment at 0μM, 0.2μM, and 0.25μM MV concentration. All values represent mean ± SD, n=2.

| **Control** | **3hr** | **6hr** | **12hr** | **24hr** | **36hr** |
|---|---|---|---|---|---|
| Normal | 93.92 ± 2.71 | 91.56 ± 3.78 | 86.14 ± 1.92 | 92.48 ± 4.74 | 92.72 ± 2.86 |
| Punctate | 6.08 ± 2.71 | 6.37 ± 3.57 | 9.65 ± 4.80 | 7.52 ± 4.74 | 7.28 ± 2.86 |
| Diffused | 0.00 ± 0.00 | 2.07 ± 0.21 | 4.21 ± 2.88 | 0.00 ± 0.00 | 0.00 ± 0.00 |
| **0.2μM MV** | **3hr** | **6hr** | **12hr** | **24hr** | **36hr** |
| Normal | 60.03 ± 1.33 | 46.88 ± 8.57 | 53.23 ± 3.00 | 83.70 ± 0.51 | 83.68 ± 5.68 |
| Punctate | 39.97 ± 1.33 | 44.96 ± 2.97 | 44.54 ± 0.14 | 12.68 ± 5.64 | 16.32 ± 5.68 |
| Diffused | 0.00 ± 0.00 | 8.16 ± 11.54 | 2.22 ± 3.14 | 3.62 ± 5.12 | 0.00 ± 0.00 |
| **0.25μM MV** | **3hr** | **6hr** | **12hr** | **24hr** | **36hr** |
| Normal | 63.10 ± 8.42 | 53.76 ± 12.77 | 51.90 ± 7.41 | 75.12 ± 6.26 | 81.67 ± 2.36 |
| Punctate | 36.90 ± 8.42 | 41.13 ± 8.83 | 44.94 ± 8.72 | 18.30 ± 3.38 | 18.33 ± 2.36 |
| Diffused | 0.00 ± 0.00 | 5.11 ± 3.94 | 3.15 ± 1.31 | 6.58 ± 2.88 | 0.00 ± 0.00 |

**Table 2.** Quantification of percentage of cells having the 3 ROS phenotypes in the chloroplast compartment at different timepoints (1hr, 3hr, 6hr, 12hr, and 24hr) upon Menadione treatment at 0μM, 20μM, and 40μM MD concentration. All values represent mean ± SD, n=2.

| **Control** | **1hr** | **3hr** | **6hr** | **12hr** | **24hr** |
|---|---|---|---|---|---|
| Normal | 95.42 ± 0.59 | 93.92 ± 2.71 | 93.41 ± 1.16 | 89.40 ± 2.69 | 92.74 ± 4.37 |
| Intermediate | 4.58 ± 0.59 | 6.08 ± 2.71 | 5.63 ± 2.52 | 7.47 ± 1.73 | 7.26 ± 4.37 |
| Punctate | 0.00 ± 0.00 | 0.00 ± 0.00 | 0.96 ± 1.36 | 3.13 ± 4.42 | 0.00 ± 0.00 |
| **20μM MD** | **1hr** | **3hr** | **6hr** | **12hr** | **24hr** |
| Normal | 89.05 ± 1.35 | 87.80 ± 1.19 | 91.29 ± 1.15 | 87.41 ± 3.95 | 89.73 ± 8.50 |
| Intermediate | 10.95 ± 1.35 | 12.20 ± 1.19 | 8.71 ± 1.15 | 12.59 ± 3.95 | 10.27 ± 8.50 |
| Punctate | 0.00 ± 0.00 | 0.00 ± 0.00 | 0.00 ± 0.00 | 0.00 ± 0.00 | 0.00 ± 0.00 |
| **40μM MD** | **1hr** | **3hr** | **6hr** | **12hr** | **24hr** |
| Normal | 80.38 ± 9.30 | 80.14 ± 8.80 | 90.28 ± 10.85 | 84.90 ± 1.15 | 89.26 ± 3.06 |
| Intermediate | 19.62 ± 9.30 | 18.73 ± 10.41 | 9.72 ± 10.85 | 15.10 ± 1.15 | 10.74 ± 3.06 |
| Punctate | 0.00 ± 0.00 | 1.14 ± 1.61 | 0.00 ± 0.00 | 0.00 ± 0.00 | 0.00 ± 0.00 |

**Table 3.** Quantification of the percentage of cells having the 3 ROS phenotypes in the mitochondrial compartment at different timepoints (1hr, 3hr, 6hr, 12hr, and 24hr) upon Menadione treatment at 0μM, 20μM, and 40μM MD concentration. All values represent mean ± SD, n=2.

| **Control** | **1hr** | **3hr** | **6hr** | **12hr** | **24hr** |
|---|---|---|---|---|---|
| Normal | 90.63 ± 5.61 | 85.95 ± 6.18 | 86.29 ± 10.91 | 80.16 ± 11.22 | 90.15 ± 7.03 |
| Intermediate | 6.04 ± 0.89 | 14.05 ± 6.18 | 13.71 ± 10.91 | 17.46 ± 14.59 | 8.63 ± 8.75 |
| Punctate | 3.33 ± 4.71 | 0.00 ± 0.00 | 0.00 ± 0.00 | 2.38 ± 3.37 | 1.22 ± 1.72 |
| **20μM MD** | **1hr** | **3hr** | **6hr** | **12hr** | **24hr** |
| Normal | 70.63 ± 2.65 | 85.43 ± 5.45 | 97.50 ± 3.54 | 97.94 ± 0.09 | 97.87 ± 3.01 |
| Intermediate | 24.69 ± 3.98 | 14.57 ± 5.45 | 2.50 ± 3.54 | 2.06 ± 0.09 | 2.13 ± 3.01 |
| Punctate | 4.69 ± 6.63 | 0.00 ± 0.00 | 0.00 ± 0.00 | 0.00 ± 0.00 | 0.00 ± 0.00 |
| **40μM MD** | **1hr** | **3hr** | **6hr** | **12hr** | **24hr** |
| Normal | 82.12 ± 5.65 | 54.86 ± 0.98 | 55.84 ± 14.16 | 89.87 ± 1.75 | 95.35 ± 0.15 |
| Intermediate | 17.88 ± 5.65 | 45.14 ± 0.98 | 43.11 ± 12.68 | 10.13 ± 1.75 | 4.65 ± 0.15 |
| Punctate | 0.00 ± 0.00 | 0.00 ± 0.00 | 1.04 ± 1.47 | 0.00 ± 0.00 | 0.00 ± 0.00 |

**Table 4.** Quantification of percentage of cells having the 3 ROS phenotypes in the mitochondrial compartment at different timepoints (3hr, 6hr, 12hr, 24hr, 36hr, and 48hr) upon Methyl viologen treatment at 0μM, 0.2μM, and 0.25μM MV concentration. All values represent mean ± SD, n=2.

| **Control** | **3hr** | **6hr** | **12hr** | **24hr** | **36hr** | **48hr** |
|---|---|---|---|---|---|---|
| Normal | 92.53 ± 7.11 | 85.59 ± 4.22 | 93.22 ± 9.59 | 90.23 ± 6.38 | 95.95 ± 5.73 | 96.94 ± 4.33 |
| Intermediate | 7.47 ± 7.11 | 14.41 ± 4.22 | 6.78 ± 9.59 | 9.77 ± 6.38 | 4.05 ± 5.73 | 3.06 ± 4.33 |
| Punctate | 0.00 ± 0.00 | 0.00 ± 0.00 | 0.00 ± 0.00 | 0.00 ± 0.00 | 0.00 ± 0.00 | 0.00 ± 0.00 |
| **0.2μM MV** | **3hr** | **6hr** | **12hr** | **24hr** | **36hr** | **48hr** |
| Normal | 91.67 ± 11.79 | 93.90 ± 5.17 | 90.95 ± 1.34 | 91.56 ± 2.50 | 89.47 ± 0.00 | 93.18 ± 9.64 |
| Intermediate | 8.33 ± 11.79 | 6.10 ± 5.17 | 9.05 ± 1.34 | 8.44 ± 2.50 | 10.53 ± 0.00 | 6.82 ± 9.64 |
| Punctate | 0.00 ± 0.00 | 0.00 ± 0.00 | 0.00 ± 0.00 | 0.00 ± 0.00 | 0.00 ± 0.00 | 0.00 ± 0.00 |
| **0.25μM MV** | **3hr** | **6hr** | **12hr** | **24hr** | **36hr** | **48hr** |
| Normal | 87.57 ± 13.85 | 90.30 ± 4.88 | 90.24 ± 2.26 | 84.27 ± 10.65 | 85.00 ± 7.07 | 94.10 ± 1.76 |
| Intermediate | 12.43 ± 13.85 | 9.70 ± 4.88 | 9.76 ± 2.26 | 4.44 ± 2.99 | 12.14 ± 3.03 | 4.11 ± 0.76 |
| Punctate | 0.00 ± 0.00 | 0.00 ± 0.00 | 0.00 ± 0.00 | 11.28 ± 13.64 | 2.86 ± 4.04 | 1.79 ± 2.53 |

**Table 5.** Quantification of the normalized ratio of Fv/Fm for all three conditions 0μM MD, 20μM MD, and 50μM MD at all the time points. All values represent mean ± SD, n=3.

| **Handy PEA** | **0.5hr** | **1hr** | **3hr** | **6hr** | **12hr** |
|---|---|---|---|---|---|
| **Control** | 0.58 ± 0.02 | 0.59 ± 0.03 | 0.60 ± 0.02 | 0.60 ± 0.01 | 0.62 ± 0.02 |
| **20μM MD** | 0.58 ± 0.00 | 0.60 ± 0.02 | 0.61 ± 0.01 | 0.62 ± 0.01 | 0.65 ±0.01 |
| **50μM MD** | 0.53 ± 0.03 | 0.46 ± 0.04 | 0.00 ± 0.00 | 0.00 ± 0.00 | 0.00 ± 0.00 |

**Table 6.** Quantification of the ratio of F (PSI/PSII) for all three conditions 0μM MD, 20μM MD, and 50μM MD at all the time points. All values represent mean ± SD, n=3.

| **77k** | **0.5hr** | **1hr** | **3hr** | **6hr** | **12hr** | **24hr** |
|---|---|---|---|---|---|---|
| **Control** | 1.11 ± 0.14 | 1.14 ± 0.15 | 1.19 ± 0.25 | 1.19 ± 0.22 | 1.15 ± 0.21 | 1.12 ± 0.17 |
| **20μM MD** | 1.02 ± 0.11 | 1.14 ± 0.22 | 1.13 ± 0.17 | 1.10 ± 0.14 | 1.22 ± 0.15 | 1.11 ± 0.17 |
| **50μM MD** | 1.16 ± 0.12 | 1.37 ± 0.18 | 0.86 ±0.30 | 0.37 ± 0.05 | 0.00 ± 0.00 | 0.00 ± 0.00 |

**Table 7.** The basal respiration **(A)**, and maximal respiration rates **(B)** were quantified for MV-treated samples at 0μM, 0.2μM, and 1μM MV across all time points. All values represent mean ± SD, n=3.

**A)**

| **Basal** | **1hr** | **3hr** | **6hr** | **24hr** | **48hr** |
|---|---|---|---|---|---|
| **Control** | 12.73 ± 6.15 | 17.67 ± 11.35 | 17.78 ± 10.44 | 12.60 ± 7.65 | 15.42 ± 4.14 |
| **0.2μM MV** | 10.62 ± 9.13 | 11.88 ± 6.96 | 10.08 ± 9.39 | 10.48 ±3.52 | 17.64 ± 2.26 |
| **1μM MV** | 15.10 ± 8.38 | 8.56 ± 5.08 | 18.71 ± 10.65 | 10.67 ± 5.06 | 3.47 ± 2.53 |

**B)**

| **Maximal** | **1hr** | **3hr** | **6hr** | **24hr** | **48hr** |
|---|---|---|---|---|---|
| **Control** | 78.77 ± 23.68 | 79.37 ± 20.26 | 86.95 ± 19.79 | 86.09 ± 17.65 | 86.42 ±15.44 |
| **0.2μM MV** | 43.45 ± 12.71 | 57.03 ± 30.33 | 61.79 ± 20.29 | 62.03 ± 26.21 | 78.32 ± 16.67 |
| **1μM MV** | 45.56 ± 34.82 | 25.65 ± 27.31 | 47.59 ± 16.69 | 29.41 ± 6.71 | 11.68 ± 5.36 |

**Table 8.** Quantification of the percentage of cells having these 3 mitochondrial morphologies at different timepoints (0hr, 1hr, 3hr, 6hr, 12hr, 24hr, 36hr, and 48hr) upon Methyl viologen treatment at 0μM, 0.2μM and 0.25μM MV concentration. All values represent mean ± SD, n=3.

| Control | 0hr | 1hr | 3hr | 6hr | 12hr | 24hr | 48hr |
|---|---|---|---|---|---|---|---|
| Tubular | 87.07 ± 2.47 | 86.32 ± 5.70 | 83.89 ± 10.29 | 81.15 ± 8.61 | 82.60 ± 5.94 | 80.19 ± 18.49 | 86.87 ± 5.47 |
| Intermediate | 6.34 ± 2.21 | 14.29 ± 0.00 | 11.00 ± 1.41 | 16.28 ± 10.65 | 10.08 ± 5.10 | 11.16 ± 6.61 | 9.17 ± 1.90 |
| Fragmented | 6.60 ±3.42 | 2.87 ± 2.53 | 11.45 ± 6.44 | 3.85 ± 2.00 | 10.98 ± 3.75 | 1.85 ± 2.62 | 5.93 ± 4.46 |
| **0.2μM MV** | **0hr** | **1hr** | **3hr** | **6hr** | **12hr** | **24hr** | **48hr** |
| Tubular | 86.75 ± 3.02 | 83.75 ± 5.30 | 95.73 ± 0.39 | 88.10 ± 6.73 | 88.38 ± 1.81 | 89.59 ± 2.25 | 90.75 ± 3.32 |
| Intermediate | 5.63 ± 2.52 | 15.00 ± 7.07 | 4.27 ± 0.39 | 8.57 ± 2.02 | 6.56 ± 4.40 | 4.94 ± 1.33 | 7.07 ± 0.25 |
| Fragmented | 7.62 ±5.54 | 2.50 ± 0.00 | 0.00 ± 0.00 | 6.67 ± 0.00 | 5.06 ± 2.60 | 5.47 ± 3.58 | 4.35 ± 0.00 |
| **0.25μM MV** | **0hr** | **1hr** | **3hr** | **6hr** | **12hr** | **24hr** | **48hr** |
| Tubular | 87.07 ± 2.47 | 81.58 ± 4.38 | 85.07 ± 7.79 | 85.10 ± 5.04 | 86.37 ± 9.18 | 78.49 ± 9.37 | 81.40 ± 6.50 |
| Intermediate | 6.34 ± 2.21 | 13.47 ± 4.24 | 12.52 ± 2.52 | 12.84 ± 4.12 | 16.52 ± 4.92 | 15.05 ± 6.54 | 10.97 ± 6.12 |
| Fragmented | 6.60 ± 3.42 | 6.60 ± 3.90 | 7.39 ± 3.41 | 4.12 ± 0.60 | 9.32 ± 7.03 | 9.36 ± 4.97 | 7.63 ± 0.64 |

**Table 9.** Quantification of the percentage of cells having different mitochondrial morphologies at specified timepoints (0hr, 1hr, 3hr, 6hr, 12hr, and 24hr) upon Menadione treatment at 0μM, 20μM, and 40μM MD concentration. All values represent mean ± SD, n=3.

| Control | 0hr | 1hr | 3hr | 6hr | 12hr | 24hr |
|---|---|---|---|---|---|---|
| Tubular | 85.65 ± 13.08 | 82.02 ± 11.95 | 85.24 ± 6.69 | 83.87 ± 6.89 | 81.19 ± 6.26 | 84.71 ± 5.64 |
| Intermediate | 14.39 ± 6.83 | 25.50 ± 1.44 | 16.03 ± 1.72 | 9.30 ± 3.31 | 11.84 ± 3.60 | 6.27 ± 5.09 |
| Fragmented | 7.12 ± 4.82 | 8.74 ± 2.26 | 3.31 ± 0.48 | 11.16 ± 3.91 | 6.97 ± 3.76 | 4.22 ± 1.41 |
| **20μM MD** | **0hr** | **1hr** | **3hr** | **6hr** | **12hr** | **24hr** |
| Tubular | 91.39 ± 7.67 | 73.11 ± 9.80 | 79.24 ± 4.67 | 85.43 ± 0.95 | 83.66 ± 5.65 | 89.99 ± 2.29 |
| Intermediate | 11.06 ± 5.16 | 23.45 ± 9.19 | 13.61 ± 4.75 | 9.52 ± 3.47 | 5.57 ± 2.22 | 4.80 ± 2.78 |
| Fragmented | 3.70 ± 0.00 | 3.44 ± 0.63 | 4.30 ± 0.65 | 3.29 ± 0.23 | 6.56 ± 1.80 | 6.50 ± 2.12 |
| **40μM MD** | **0hr** | **1hr** | **3hr** | **6hr** | **12hr** | **24hr** |
| Tubular | 85.65 ± 13.08 | 75.36 ± 14.96 | 60.65 ± 9.16 | 64.46 ± 13.97 | 78.87 ± 7.12 | 79.67 ± 6.96 |
| Intermediate | 10.79 ± 9.10 | 15.90 ± 3.25 | 28.50 ± 4.34 | 11.80 ± 1.57 | 13.14 ± 4.93 | 13.93 ± 4.74 |
| Fragmented | 7.12 ± 4.82 | 17.48 ± 16.87 | 14.46 ± 2.15 | 15.97 ± 12.35 | 10.50 ± 6.27 | 21.74 ± 0.00 |
